# Archerfish number discrimination

**DOI:** 10.1101/2021.10.04.463045

**Authors:** Davide Potrich, Mirko Zanon, Giorgio Vallortigara

## Abstract

Debates have arisen as to whether non-human animals actually can learn astract non-symbolic numerousness or whether they always rely on some continuous physical aspect of the stimuli covarying with number. Here we investigated archerfish (*Toxotes jaculatrix*) non-symbolic numerical discrimination with accurate control for co-varying continuous physical stimulus attributes. Archerfish were trained to select one of two groups of black dots (Exp. 1: 3 *vs*. 6 elements; Exp. 2: 2 *vs*. 3 elements); these were controlled for several combinations of physical variables (elements’ size, overall area, overall perimeter, density and sparsity), ensuring that only numerical information was available. Generalization tests with novel numerical comparisons (2 *vs*. 3, 5 *vs*. 8 and 6 *vs*. 9 in Exp. 1; 3 *vs*. 4, 3 *vs*. 6 in Exp. 2) revealed choice for the largest or smallest numerical group according to the relative number that was rewarded at training. None of the continuous physical variables, including spatial frequency, were affecting archerfish performance. Results provide evidence of the spontaneous use of abstract relative numerical information in archerfish for both small and large numbers.

## INTRODUCTION

Non-symbolic numerical estimation is an important and well-studied cognitive ability that allows humans and other animals to interact successfully with their surroundings. The development of a “sense of number” is associated with fundamental biological needs that in many ecological contexts allow animals to estimate how many companions or enemies are around, or how much food is present in different patches - all important information to maximize fitness and reproductive success in the wild [1].

Typically, in order to assess numerical abilities animals are requested to discriminate between sets of visual stimuli differing in numerosity (review in [2]). This can be done using spontaneous attractive natural stimuli such as food or social companion, taking advantage of the animals’ natural and spontaneous tendency in some ecological contexts to “go for more”. Alternatively, operant conditioning procedures can be used that associate a particular set of stimuli with a reward. Extensive evidence supports the use of numerical information in non-human primates (e.g., [3–7]), as well as in other mammals (e.g., [8–13]), in birds (e.g., [14–19]), in amphibians (e.g., [20,21]), in reptiles (e.g., [22,23]), in fish (e.g., [24–26]) and in arthropods (e.g., [27–30]) (see for general reviews in vertebrates [1,31,32]).

Numerical discrimination seems to be supported by an “Approximate Number System” (ANS, [33,34]), which discriminative accuracy is ratio-dependent in accordance to Weber’s law (as the ratio between two numerosity increases, the discrimination gets more difficult). Besides the ANS, an attentional working memory-based system has been claimed for by some authors as providing precise representation of small numbers (up to 3-4), the so-called “Object Tracking System” (OTS; [35]), though its generality for non-human animals is debated (discussion in [31,36]).

Studies investigating the neural basis of number representation revealed selectivity of response of neurons in some areas of the brain such as the parietal and prefrontal cortex in humans [37,38] and in monkeys [39,40], the nidopallium caudolaterale in crows [15,41] and the most caudal dorsal-central part of the pallium in zebrafish [42,43], suggesting that common selective pressures led to convergent evolution of numerical representation in different species [44,45].

However, one issue in all these experiments is that animals are dealing with sets of physical elements, and thus numerical information is intrinsically melted with other non-numerical properties of the stimulus, such as the area, the density or the spatial frequency or the elements’ arrangement [46]. Recently, some debates have arisen concerning whether bees use abstract numerical information or rather rely on sensory properties of the stimulus for discrimination [47,48].

Taking advantage of the fact that we recently developed a sophisticated script for the automatic generation of visual stimuli that can allow proper randomization and control of continuous physical variables in number sense experiments [49], we decided to perform some very precisely controlled experiments to check whether fish do use number as abstract property.

We selected archerfish (*Toxotes jaculatrix*) for our study. These fish are well-known for their particular hunting strategy, which consists of spitting at preys above the water surface with a precise jet of water thrown with the mouth. This attacking repertoire makes it very easy to train them to hit targets using operant conditioning (see e.g., [50]).

Still, to date, no studies in archerfish have explicitly investigated abstract numerical abilities. Leibovich-Raveh et al. [51] and colleagues showed that when archerfish make magnitude-related decisions, their choice is influenced by the non-numerical variables that positively correlate with numerosity; for instance, when exposed to two groups of dots differing in number and continuous physical information, archerfish spontaneously selected the group containing the larger non-numerical magnitudes and smaller numerosity, switching to the larger numerical set when positively correlated with all the non-numerical magnitudes.

Related to magnitude discrimination, archerfish also proved to be able to associate different geometric shapes with different food quantities [52]; this would support the existence of a system dealing with magnitudes, although a specific role of numerical information remains unclear.

In our study archerfish were trained to select one of two arrays, involving either a small and a large numerosity (Exp. 1: 3 *vs*. 6 elements) or small numerosities only (Exp. 2: 2 *vs*. 3 elements). After reaching a learning criterion, archerfish were tested with novel numerical comparison (2 *vs*. 3, 5 *vs*. 8 and 6 *vs*. 9 in Exp 1; 3 *vs*. 4, 3 *vs*. 6 in Exp 2) to check whether the rule they used in the training phase was based on a relative judgement (select the “largest” or “smallest” group) or on an absolute judgment (select a specific number of item). All of the different continuous physical variables such as radius, total area, total perimeter, convex hull and inter-distance were carefully controlled for and alternately balanced across trials, ensuring that the animals could not rely on them to perform their judgment (Figure 1). Furthermore, a statistical analysis was run for *a posteriori* evaluation of whether any of these variables influence the archerfish responses, confirming that they were not used as a cue for numerical evaluation.

**Figure 1.**
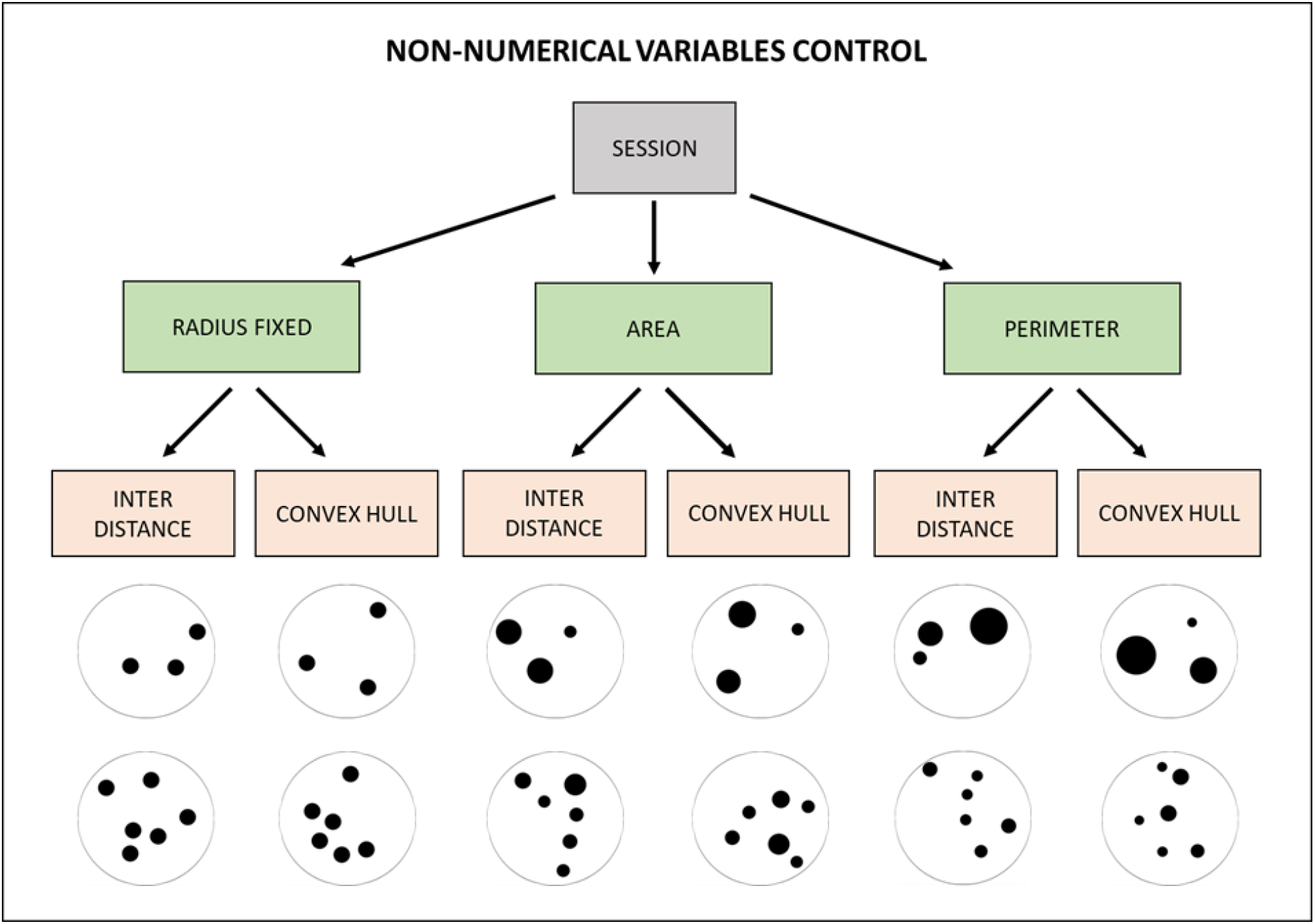
Schematic representation of the non-numerical physical controls applied to the stimuli in each session.

## RESULTS

### EXPERIMENT 1

Eight archerfish were trained to discriminate between two groups of black dots in a 3 *vs*. 6 numerical comparison; four fish were trained to select the number 3, while the other four were rewarded with the number 6. No difference has been found in the number of trials needed to reach the learning criterion between the group trained with 3 elements (mean±SEM= 451.25±106.77) and the group trained with 6 elements (mean±SEM= 413.25±73.14) (Independent Samples t-Test: t(6)=0.294, *p*=0.779).

Once the learning criterion was reached, all the fish performed three different tests.

#### TEST 1

This test was the main discriminator to understand whether at training fish represented numerosity as relative or absolute. Fish trained to select the smallest number 3 at training (i.e. the smallest set in the 3 *vs*. 6) were presented at test with a novel discrimination 2 *vs*.3, while fish trained to select the number 6 at training (i.e., largest set in the 3 *vs*. 6) were tested with a 6 *vs*. 9 condition. The use of “relative” information (go for the smallest or largest) should lead the fish to choose the novel numerosity at test, while the use of “absolute” information would reflect in the choice of the stimulus with the same number of elements as at training.

#### TEST 2

The second test aimed to clarify the role of the incorrect (i.e., unrewarded) training stimulus and its relevance for the fish. When fish are trained to select the numerosity 3, thus avoiding number 6, once presented with the new comparison 6 *vs*. 9 (or *vice versa* 2 *vs*. 3, if trained to select 6), do they choose the group according to the relative information even if it coincides with the absolute numerosity to avoid at training?

#### TEST 3

In the last test, fish behaviour was observed in a comparison involving novel numerosities never experienced during the training, i.e., 5 *vs*. 8. This allowed observing whether zebrafish applied a relative representation (go for the “smallest” or “largest”), or if the choice was at the chance level, since no absolute numerical information experienced at training was present here.

Results at tests for Experiment 1 are reported in Figure 2. Choices for the relative numerosity were analyzed using a generalized linear mixed model fit by maximum likelihood (Laplace Approximation), binomial GLMM with a logit link in R. Four fixed effects (type of Training -3 or 6 dots-; type of Test -2 *vs*. 3, 5 *vs*. 8 and 6 *vs*. 9-; type of geometrical control -radius fixed, overall area controlled, overall perimeter controlled-; type of spatial disposition control -inter-distance controlled; convex-hull controlled-) and one random intercept effect (fish) were considered. Analysis on the random effect showed not to affect the model and no significant differences were found between effects of groups, nor group interactions (comparisons between different models considering various effects and interactions reported always p> 0.05, suggesting to adopt the simplest model described by the only choice with no contribution from any effects). Only a trend for the contribution of the type of geometrical control was observed, driven by a non-significant difference between the “radius fixed” and “overall area controlled” conditions (post-hoc non-parametric tests adjusted with Tukey method: p = 0.063); within this trend, every single condition was statistically significant by chance level in the direction of the relative choice (“radius fixed”: probability of success = 71.18%, p < 0.001; “overall area controlled”: probability of success = 81.25%, p < 0.001; “overall perimeter controlled”: probability of success = 73.61%, p < 0.001).

**Figure 2.**
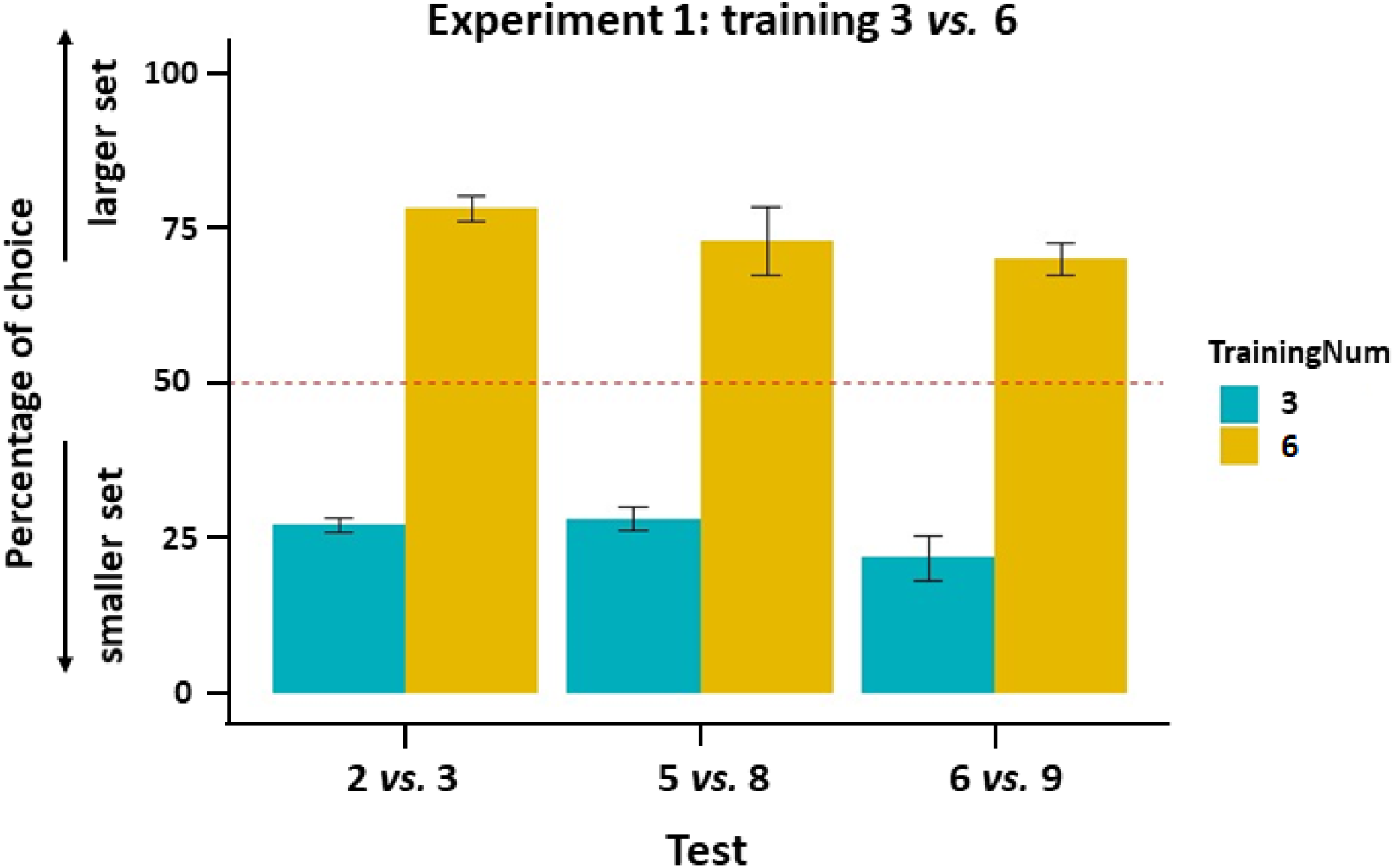
Percentage of choice for the larger/smaller set (mean ± SEM) displayed by fish in the novel comparison tests for the groups trained to select the smaller (3) or larger (6) set.

Overall, considering the previous discussion, a binomial test shrinking all the data together was performed to investigate the final findings: fish showed a strong significant preference for the relative numerosity (probability of success: 74.3%, p < 0.001).

The result obtained in Experiment 1 showed that archerfish, when trained to select one of two simultaneously displayed groups of dots with different numerosities (i.e., 3 *vs*. 6 dots), use a relative numerical rule to perform novel numerical comparisons. These results confirm findings in other fish species such as angelfish [55] and guppy [56] but they are different from those obtained in bees which showed instead a preference for the absolute number [57]. An important difference between fish and bees studies is related to the numerical comparison used: respectively large numbers (> 4 elements) for fish and small numbers (≤ 4 elements) with bees. This might engage different systems (see Introduction) thus explaining the discrepancy. The training discrimination used here in Experiment 1 involved two numbers (3 *vs*. 6) that belong one to the hypothesized “small” and the other to “large” systems, respectively. This is different than in previous fish studies which involved only large numerosities; thus, it remains to be tested how fish would deal when trained with small numerosities only. In principle, the presence of a large number in the comparison in Exp. 1 may be enough to lead the archerfish to follow a relative rule. If trained with a numerical discrimination involving only small numbers, would the animals still use a relative numerosity judgement or would they turn to absolute judgement? This was tested in Experiment 2.

### EXPERIMENT 2

Four subjects were trained to select the largest number in a 2 *vs*. 3 comparison (i.e., the number 3). Fish judgment was then observed in two tests (i.e. 3 *vs*. 4 and 3 *vs*. 6) involving a comparison between the previously trained numerosity (3) and a novel numerosity (4 or 6).

All fish reached the learning criterion, showing an ability to discriminate between the two numbers (trials to criterion±SEM= 506.5 ± 97.8). Results at test are reported in Figure 3. A GLMM model with three fixed effects (type of test -3 *vs*. 4 and 3 *vs*. 6-; type of geometrical control -radius fixed, overall area controlled, overall perimeter controlled-; type of spatial disposition control -inter-distance controlled; convex-hull controlled-) and one random intercept effect (fish) showed no random effect of fish, neither significant differences between groups or groups’ interactions (Chi-Square tests between all different models with different effects and interactions report always p> 0.05, suggesting to adopt the simplest model based on the only fish choice -for the relative or absolute number- and no effects of controls).

**Figure 3.**
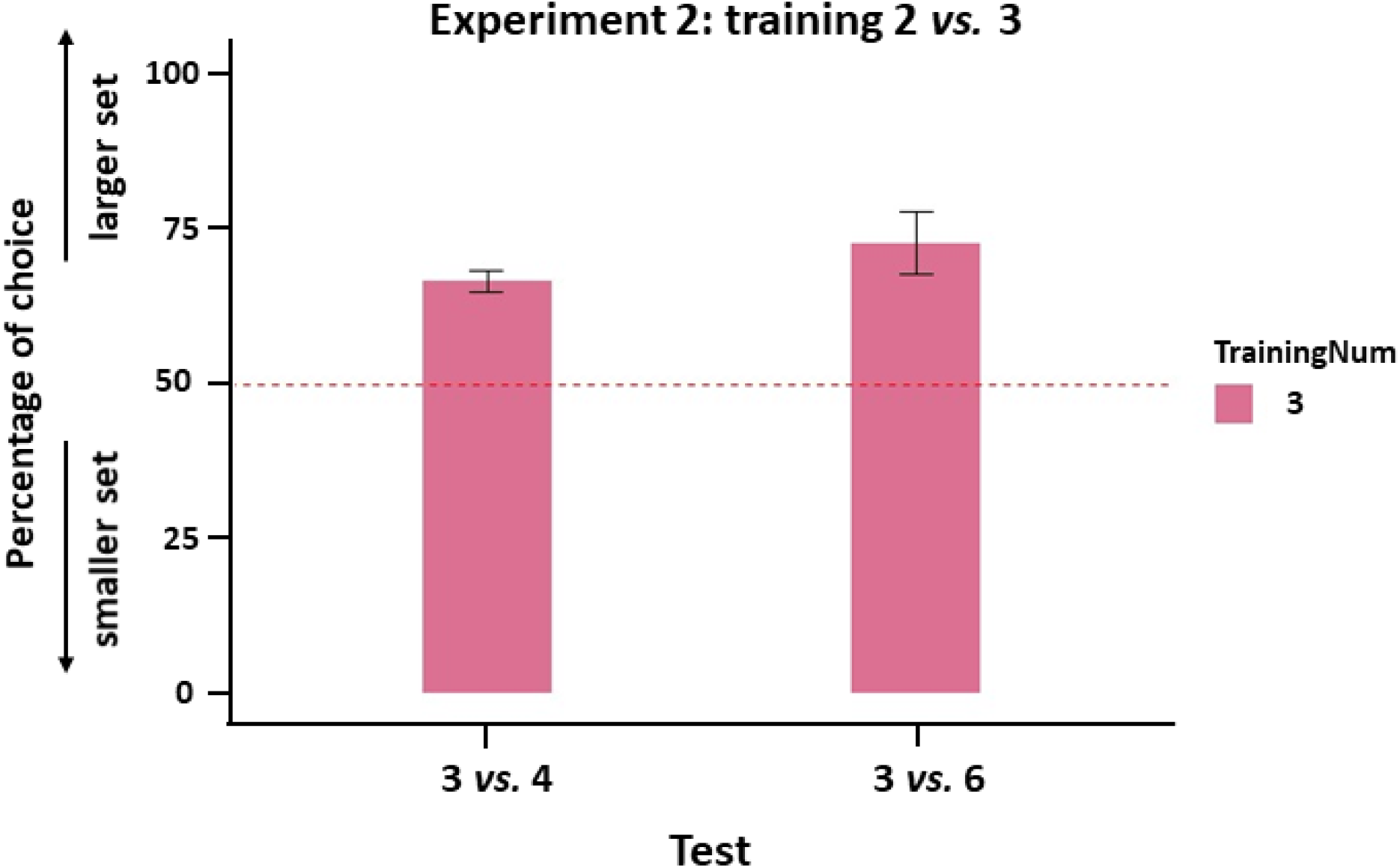
Percentage of choice for the larger/smaller set (mean ± SEM) displayed by fish in the novel comparison tests for the group trained to select the larger (3) set.

An Exact binomial test considering a merge of the data showed a highly significant preference for the relative number (probability of success: 69.79%, p < 0.001).

In Experiment 2, archerfish showed to be able to discriminate between two different numerical groups of dots within the small numerical range. At test, fish preferred the novel numerosity to the familiar 3 items, in both 3 *vs*. 4 and 3 *vs*. 6 comparisons, confirming the use of a relative rather than absolute numerical rule. This evidence does not match with findings in bees, tested in the same numerical conditions, suggesting that the spontaneous engagement of relative/absolute rule to extract numerical information may be guided by different ecological pressures experienced by different species in their phylogenetic history. The spontaneous use of relative rules suggests that among fish, it is more important to learn a general rule that is applicable to novel comparisons. It cannot be excluded that this strategy is adopted because it could be less demanding as to memory load than an absolute judgement strategy.

Considering the results of Experiment 2 and Experiment 1, it is apparent that archerfish can easily discriminate between small and large numerosity using the same rules, providing evidence in favour of a unique system underlying numerical discrimination as found in other fish species [26,58].

### NUMEROSITY AND SPATIAL FREQUENCY

The stimuli used in our Experiments were visual collections of black dots differing in numerosity. As described in the method section, for each numerical comparison, the physical properties of each array were equalized for the geometry (radius, area and perimeter) and spatial disposition (inter-distance (density) and convex-hull; see Figure 1). Since we are dealing with images, each figure could also be described in terms of spatial frequency. Spatial frequency can be thought of as the number of repeating elements in a pattern per unit distance, and it is mathematically described by the Fourier transform theory. No control was applied to the spatial frequency of our stimuli. Thus, in order to check whether spatial frequency could influence archerfish choice, we calculated spatial frequency variation across all different numerosities and control conditions (see Methods section). Within each numerical test comparison, different spatial frequencies were found (see Figure 4). The different constraints applied to the stimuli (control of the area, perimeter or elements radius) showed to influence differently the spatial frequency between the two numerosities. In detail, when the elements’ radius was fixed between the two numerical arrays, the total power of the spatial frequency was higher in the smaller group than in the larger one, while the opposite was found in the groups in which the overall perimeter was balanced (total power higher in the more numerous group). Interestingly, this trend was maintained in all the numerical comparisons used, irrespective of the number of elements to be compared.

**Figure 4.**
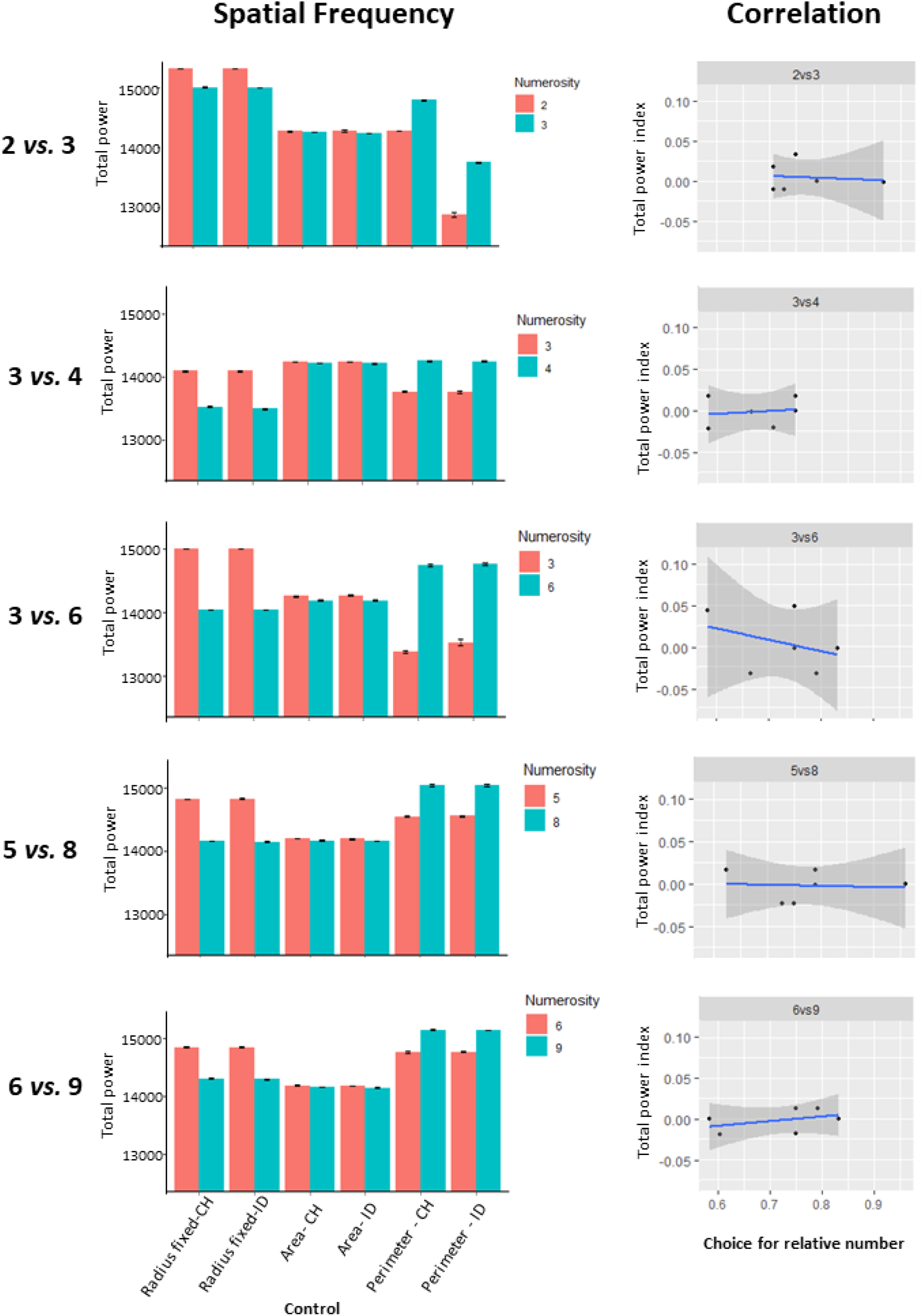
The histograms (on the left) show the spatial frequency (Total power) for each numerical comparison among the different control groups (non-numerical variables control). The different constraints applied to the stimuli (control of the area, perimeter or elements radius) showed to influence the spatial frequency between the two compared numerosities. The regression lines (on the right) show the correlation between fish’ performance accuracy (choice for the relative numerosity) and the spatial frequency (total power index between the two total power values), for all numerical comparisons.

To investigate the influence of spatial frequency in the numerical task, we analyzed whether a correlation between the performance accuracy and the spatial frequency was apparent, for all possible control configurations (see Methods section). Results are reported in Figure 4, showing no correlations between any comparison (test 2 *vs*. 3: r(4)= -0.17, p=0.83; test 3 *vs*. 4: r(4)= 0.15, p=0.77; test 3 *vs*. 6: r(4)= -0.35, p=0.50; test 5 *vs*. 8: r(4)= -0.08, p=0.88; test 6 *vs*. 9: r(4)=-0.42, p=0.41. These data strongly suggest that the spatial frequency was not influencing archerfish performance in the numerical task.

## DISCUSSION

Overall, our results showed that when trained to select a specific group of elements between two numerical arrays, archerfish spontaneously generalize at test to novel numerical comparison according to a relative numerical rule (select the largest/smallest) rather than an absolute numerical rule (select the specific number of items). These findings are in agreement with previous results from other fish species and humans [55,56], while differing with respect to bees [57].

Interestingly, archerfish uses a general relative judgement even when trained to discriminate between numerosities that belong to different systems, namely small and large numerosities (for a review see [61]). In Experiment 1, archerfish were trained with a 3 *vs*. 6 contrast and then observed in test conditions with a 2 *vs*. 3, 6 *vs* .9 and 5 *vs*. 8 comparison. In all the tests, archerfish showed to spontaneously use a general relative rule. In Experiment 2, subjects’ performance was observed in a numerical discrimination involving at training only small numerosities (i.e., 2 *vs*. 3). Once again, at test, fish followed the relative rule, selecting the largest group in the test comparisons 3 *vs*. 4 and 3 *vs*. 6, thus ignoring the absolute number of elements (3).

Taken together, our results support the hypothesis of a unique system for representing numerosities in archerfish, working both for small and large numbers, obeying the ANS. Evidence from other fish species supports this claim [26,58].

The reason for which archerfish primarily rely on the relative information of numerical groups may have ecological reasons, being more adaptive in a natural environment that constantly require numerical/quantity judgement. Selecting the largest social group of companions or the largest food patch are easy rules that can be more efficient than using an absolute rule. Moreover, the use of relative information may be less cognitively expensive (in terms of memory load) than the absolute one, since it does not require storing the information about the precise number of elements: the discrimination could work on a simple relative comparison between numerosities, guided by the numerical ratio between the two. Nevertheless, the engagement of relative rules requires a good level of abstraction and the creation of a general rule to be applied to [62].

In fish, the use of an absolute rule may not be as convenient as the relative one, given that in most ecological contests there is not a specific optimal amount of food, partners or companions. However, this seems not to be the case for species such as bees, which showed instead a spontaneous use of absolute numerical information, suggesting that this rule may be more informative and useful in their ecological environment. Similar evidence has been found in spiders, that, in a natural predatory strategy context, settle their attack based on the specific number of conspecifics at the nest [29].

Note, however, that the spontaneous use of a relative or absolute rule does not imply that animals are unable to use both. Vertebrates can be trained to learn a specific number of items in a set if forced to do it [6,56,63,64]. Similarly, bees can be trained to the numerical concepts of “greater than” or “smaller than” [47]. The spontaneous engagement of one of the two criteria is therefore justified probably by a combination of natural constraints and/or less cognitive demand motivations that better fit for the individuals’ fitness in their particular niches of adaptation.

Lastly, with respect to the main question of our paper, the results showed that archerfish are capable of abstract numerical discrimination, not influenced by other continuous physical variables. We tested archerfish with numerical arrays well controlled for all the possible non-numerical variables (e.g., total area, perimeter, inter-distance (density), convex hull), thus ensuring that the discrimination made by the animals was based on purely numerical information. The results of the statistical analyses showed no influence whatsoever of the different control conditions on the fish choices. Moreover, we showed that even the different spatial frequencies of the stimuli were not influential on archerfish performance. The total power of the spatial frequency has been described in the literature to positively increase with numerosity [48]; however, in our stimuli, the different geometrical constraints showed that it can be reversed as well. Moreover, elements area and perimeter seem to play a crucial role in the distribution of the spatial frequencies’ energy with respect to the elements disposition (inter-distance and sparsity). All our analyses suggested that the amplitude component of the spatial frequency was not influencing archerfish numerical evaluation during our experiments.

Note, however, that in all studies carried out so far (including our own analysis), the focus was on the amplitude of the spatial frequency as the main component, which provides information on the alternation rate of different elements in the image. It is likely that a more specific role on computation of numerosity is played by the spatial frequency phase component (related to elements’ spatial coherence and distribution) which directly relates to figure-ground aggregation and unity formation.

In conclusion, our results provide clear evidence that under conditions of strict control of continuous physical variables archerfish can encode an abstract concept of number to support relative numerical judgement for both small and large numerosities.

## MATERIALS AND METHODS

### SUBJECTS AND REARING CONDITIONS

Sixteen adult archerfish, *Toxotex jaculatrix* (fish size ranged between 8 and 10 cm in length) were provided by a local commercial supplier (“Acquario G di Segatta Stefano”). Four animals were excluded because they did not show any consistent motivation in hitting the screen. A group of fish (N=8) took part in Experiment 1, while a second group (N=4) took part in Experiment 2. All fish were housed in large aquariums (100 × 40 × 40 cm) in groups of 10 individuals. Prior to the experiment, each archerfish was moved into individual aquaria (40 × 30 × 50 cm) filled with freshwater maintained at 25°C and enriched with gravel and a shelter. Water quality was kept by suitable filters (Sera fil 60). The system was illuminated under a 10:14 light /dark cycle (Sylvania luxline plus F36W/840 cool white). Fish were fed with food pellets (Hikari cichlid gold baby pellet).

### APPARATUS

Both the apparatus and the training method were set up based on previous studies conducted with archerfish on visual discrimination tasks (i.e., [52,53]). Each experimental tank consisted of a rectangular aquarium with a monitor screen located above it (20”, DELL 2009Wt), held at 30 cm from the water level (Figure 5a). Each tank was surrounded by white opaque panels to ensure that the fish was not distracted by external cues. Each tank was raised 8 cm off the table thanks to lateral supports, allowing the positioning of a video camera under the centre of the pavement’s tank to record a bottom view of the fish and the screen (see video example in the supplementary materials).

**Figure 5.**
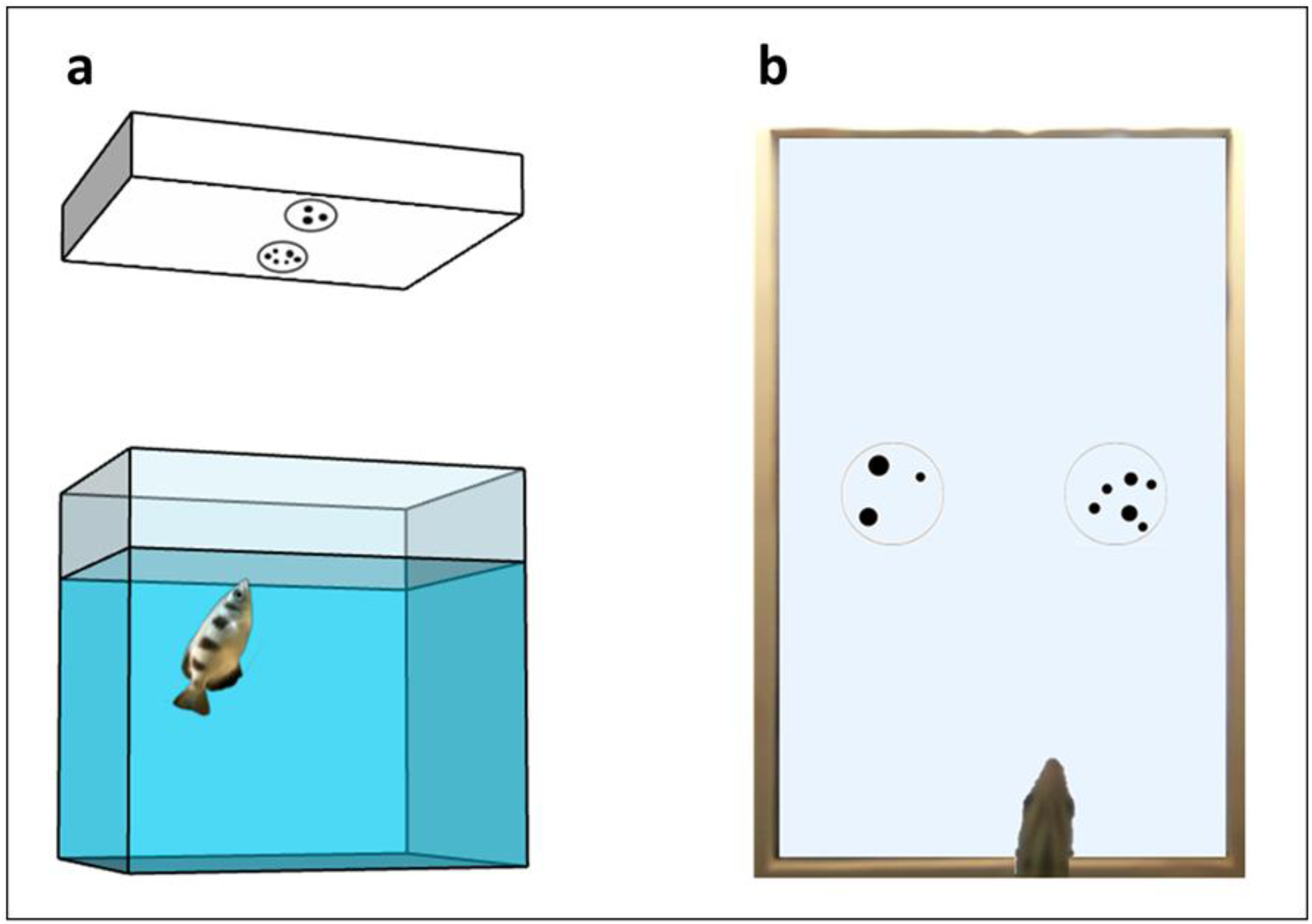
a) Schematic representation of the experimental setup; b) Bottom view of the tank from the camera placed below the tank’s pavement.

### STIMULI

The stimuli presented in the training phase consisted of groups of black dots confined into a black outline circle (6 cm diameter). The dots size was ranging between 3 and 12 mm, and the visual angle was in the range 0.43° and 1.72°, which has been proven to be well perceived by archerfish [54]. In every trial, a couple of stimuli was simultaneously presented in the centre of the screen (horizontally aligned to the shortest monitor’s side, see Figure 5b). All the stimuli were created using the software GeNEsIS [49], a Matlab program that allows to create numerical collections of stimuli controlled for several non-numerical magnitudes. Given that it is mathematically impossible to balance all the non-numerical magnitudes simultaneously in two different numerical groups, different sets of stimuli were created for each numerosity, controlling for some visual physical property; all the possible properties were covered across the different sets during a session (see Figure 1 for a view of all the combinations applied in a session). Pictures from each set were randomly presented, making the numerical information the only reliable cue to differentiate the two stimuli across all the various trials.

### GENERAL PROCEDURE

#### Pre-training phase

Before starting the experiment, fish underwent a pre-training phase in which they were gradually habituated to spit (hit with a jet of water) at the training stimulus on the screen. This was accomplished throughout a shaping procedure to facilitate the task. The silhouette of an insect was initially presented, inducing the fish reaction to spit at the prey; once hit, fish were rewarded with a food pellet. The insect was gradually replaced by a black dot and finally with the effective training stimulus. Once the fish accomplished all these stages, the training phase was initiated.

#### Training phase

Fish were trained to spit at the correct target presented on the monitor above the tank. The stimuli to discriminate consisted of two groups of dots with different numerosity. Every trial started with the appearance of a blinking black square (1.6 cm, three blinks of 100 milliseconds) at the centre of the screen to catch the fish’s attention towards the screen. Then, the two training stimuli were displayed one next to the other (distance 7 cm) on the two sides of the monitor. Only one of the two numerosities was rewarded with a food pellet when hit, while the choice for the incorrect stimulus caused the stop of the trial, which in every case, in absence of choice, was stopped after 5 minutes. At the end of each trial, the screen was cleaned from the water drops and a new trial started. In the first training session only, a corrective method was applied: the stimuli remained on the screen until the subject selected the correct target, even if the incorrect stimulus was hit, allowing the fish to correct its choice.

Fish were trained with daily sessions of 48 trials, in which continuous physical variables were controlled and changed according to the scheme reported in Figure 1, and the position of the target stimulus on the screen (right-left) was randomized. Fish generally responded 70% to 100% of the trials. The learning phase was considered completed when the fish reached a learning criterion of at least 75% of correct choices for two consecutive days (binomial test: p < 0.01), allowing the fish to take part in the test phase.

#### Test phase

Generally, each test condition consisted of the presentation of a couple of stimuli with a novel numerical comparison, aiming to see if the numerosity target learned in the training phase was represented as a relative or an absolute numerical information. Each test was composed of 24 probe trials not rewarded, divided into three testing days of 8 trials. In each test session, the 8 test trials were shuffled and interspersed with rewarded recall training trials (32 recall in total), to maintain the fish motivation high during the whole test duration. The order of the tests was randomized among the fish to exclude that the performance could be influenced by their order. At the end of each test, the fish underwent a complete daily session of retraining to further exclude potential interference among the tests.

#### Statistical analyses and data analysis

Data were analyzed using R software (R-4.1.0). In Experiment 1, an independent t-test was used to compare the number of trials to reach the criterion between the two groups at training. At test (Exps. 1 and 2), choices for the relative numerosity were analyzed using a generalized linear mixed model fit by maximum likelihood (Laplace Approximation), binomial GLMM with a logit link. A binomial test was used to compare the distribution of the choices for the relative and absolute numerosities.

To obtain an estimate of the spatial frequency we adopted an approach already performed in other studies [48,59,60]: the fast Fourier transform of our images was calculated, a radial average of the signal amplitude in the frequency domain was performed, and lastly, all the frequency contributions of its power spectrum were summed up. In this way, a value related to the total energy of each frequency component inside a given image is obtained.

To investigate the influence of spatial frequency in the numerical task, we analyzed whether a correlation between the performance accuracy (choice for the relative numerosity) and the spatial frequency (normalized total power difference between the two compared numerosities) was apparent, for all possible control configurations. To compare two numerosities we reported a normalized difference (total power index) between the two total power values (difference between the total power of the biggest numerosity and the smallest, divided by their sum). All the frequency calculations were performed with a custom script in Matlab, while the statistical comparisons were calculated in R. For each of them a Pearson’s correlation coefficient was calculated comparing the choice for the relative numerosity and the normalized difference between numerosities (as explained above).

## Ethical regulation

The present research was carried out at the Animal Cognition and Neuroscience Laboratory (ACN Lab) of the CIMeC (Center for Mind/Brain Sciences), at the University of Trento (Italy). All husbandry and experimental procedures complied with European Legislation for the Protection of Animals used for Scientific Purposes (Directive 2010/63/EU) and were approved by the Scientific Committee on Animal Health and Animal Welfare (Organismo Preposto al Benessere Animale, OPBA) of the University of Trento and by the Italian Ministry of Health (Protocol n. 932/2020-PR).

## Authors’ contribution

D.P. and G.V conceived the study. D.P. G.V and M.Z. designed the experiment. D.P. performed the experiments. All authors interpreted the data and contributed to the manuscript writing.

